# Comparison of trimethoprim-sulfamethoxazole versus ciprofloxacin monotherapy in *Stenotrophomonas maltophilia* bloodstream infections in children

**DOI:** 10.1101/339291

**Authors:** Ahu Aksay, İlker Devrim, Nagehan Katipoğlu, Nuri Bayram, Sertaç Arslanoglu, Şebnem Çallkavur, Gamze Gülfidan, Gökhan Ceylan, Hasan Ağın

## Abstract

Treatment of invasive infections caused by *Stenotothromhomas maltophilia* is difficult as the bacterium is frequently resistant to a wide range of commonly used antimicrobials. The aim of this retrospective study was to evaluate effectiveness of treatment with ciprofloxacin monotherapy compared to trimethoprim-sulfamethoxazole monotherapy in patients with *S. maltophilia* infections. We evaluated 50 patients with *S. maltophilia* bloodstream infections who had received monotherapy with trimethoprim-sulfamethoxazole or ciprofloxacin. Twenty-five patients (50%) received trimethoprim-sulfamethoxazole, and 25 patients (50%) received ciprofloxacin. Thirty-six (72%) patients were in the intensive care unit. All of the patients had at least one indwelling devices and approximately 25% patients had immunosuppression. Ciprofloxacin had the same clinical outcome and mortality as trimethoprim-sulfamethoxazole for the treatment of *S. maltophilia* bacteremia. In our study; side-effects ratio were not statistically different between ciprofloxacin and trimethoprim-sulfamethoxazole group. Ciprofloxacin could be an alternative drug of choice for treating *S. maltophilia* infections especially in premature infans in stead of trimethoprim-sulfamethoxazole.

## INTRODUCTION

*Stenotrophomonas maltophilia* is a Gram-negative obligate aerobe that is rod shaped and motile with a few polar flagella (1). In severely ill patients, *S. maltophilia* causes a wide range of infections such as bacteremia, pulmonary infections, urinary tract infections, wound infections, meningitis, peritonitis, ocular infections and endocarditis (2–4). The major predisposing factor for *S*. *maltophilia* infection in hospitalized patients is the implantation of medical devices such as central venous catheters (5), urinary tract catheters (6), prosthetic heart valves (7), endotracheal tubes (8) and intraocular (9) and contact (10) lenses. Additional risk factors for *S. maltophilia* infections include immunosuppression, neutropenia, recent broad-spectrum antibiotic therapy, including carbapenems; and prolonged hospital stay (9). *S. maltophilia* is not a highly virulent pathogen, but it has emerged as an important nosocomial pathogen associated with crude mortality rates ranging from 14 to 69% in patients with bacteremia (11, 12). Treatment of invasive infections caused by this organism is difficult as the bacterium is frequently resistant to a wide range of commonly used antimicrobials. Antibiotics with *in vitro* activity against *S. maltophilia* include trimethoprim-sulfamethoxazole (TMP-SMX), fluoroquinolones (FQs), tetracyclines, ticarcillin-clavulanate, and ceftazidime; however, there are limited clinical data focusing on *S*. *maltophilia* infections in children (5,13–17). TMP-SMX is recommended as the agent of primary choice for the treatment of *S. maltophilia* infections, but FQs are an attractive option due to *in vitro* activity (18). The purpose of this study was to evaluate patients with *S. maltophilia* infections and compare the treatment regiments of ciprofloxacin (CIP) monotherapy compared to TMP-SMX monotherapy.

## MATERIAL AND METHODS

This retrospective study included patients with *S. maltophilia* bloodstream infections who had received monotherapy with TMP-SMX or CIP. The study population included patients with positive blood cultures for *S. maltophilia* who were younger than 18 years old. The bloodstream infections were diagnosed according to CDC (19). The patients who received treatment for *S. maltophilia* for at least 48 h were included in the study. Patients were excluded if they received combination therapy for *S. maltophilia*. All patients with positive blood cultures for *S. maltophilia* were identified from microbiological reports from January 2014 to February 2015. Identification and determination of antibiotic susceptibility were performed with the VITEK-2 system (BioMerieux, France). Antibiotic susceptibility testing was done as per Clinical and Laboratory Standards Institute (CLSI) guidelines. The dosage of CIP was 20-30 mg/kg/day in 2 divided doses (max dose: 1.5 g/day) and TMP-SMX was 20 mg TMP/kg/day divided every 6 hours.

### Data

Demographic information was collected from the patients’ electronic medical records, including underlying illnesses, presence of indwelling devices, immunosuppression, and prior antibiotic use. Microbiological cure at the end of therapy (EOT), clinical response at EOT, in-hospital and 30-day mortality, and isolation of a nonsusceptible isolate within 30 days of EOT were recorded. The clinical response at EOT was evaluated and determined by improvement in all signs and symptoms of infection with no further treatment required. Microbiological cure was defined as a negative culture, from the same site as the original positive culture, at or prior to the EOT. Patients who did not have consecutive culture were excluded from analysis for this endpoint.

### Statistical analysis

Categorical variables were analyzed using a chi-square or Fisher exact test. Continuous variables were analyzed using Student’s *t* test or the Mann-Whitney U test. A *P* value of <0.05 denoted statistical significance. Data were analyzed using SPSS software version 20.0 (IBM Corp., Somers, NY).

The study was approved by local ethics committee of Dr. Behçet Uz Children’s Hospital.

## RESULTS

A total of 50 patients were included in this study with a median age of 3 months (ranging from 3 days to 17 years). Among them, 29 patients (58%) were female and 21 patients (42%) were male. Thirty-six (72%) patients were in the intensive care unit (ICU) including neonatal intensive care unit (22 [44%]), pediatric intensive care unit (9 [18%]), and cardiovascular intensive care unit (5 [10%]) at the time of culture positivity. Other patients were in the pediatric infectious disease unit (11 [22%]), pediatric hematology-oncology department (2 [4%]), and burn unit (1 [2%]). Twelve of 22 patients (54.5%) were premature infants hospitalized in the neonatal intensive care unit.

Forty patients (80%) had bloodstream infections and 10 patients (20%) had catheter-related bloodstream infections with *S. maltophilia*. Twenty-five patients (50%) received TMP-SMX, and 25 patients (50%) received CIP. The most common underlying co-morbid conditions were neurometabolic diseases (38%), leukemia (22%), cardiac diseases (8%), respiratory distress syndrome (6%), necrotizing enterocolitis (4%), and nephrotic syndrome (4%). The most common type of indwelling device was endotracheal tube (58%), followed by a central venous catheter (CVC) (30%) and genitourinary catheter (24%). Pulmonary infections accounted for 24% of all *S. maltophilia* infections. There were 4 (8%) pulmonary infections in patients who received TMP-SMX and 8 (16%) in patients who received CIP. Seventeen (34%) patients received an antibiotic prior to isolation of *S. maltophilia.* Cephalosporins (18%) and carbapenems (16%) were the most common used antibiotics (Table 1). No difference was present between TMP-SMX and CIP group by means of demographic features (p>0.05) (Table 1).

**Table-1:**
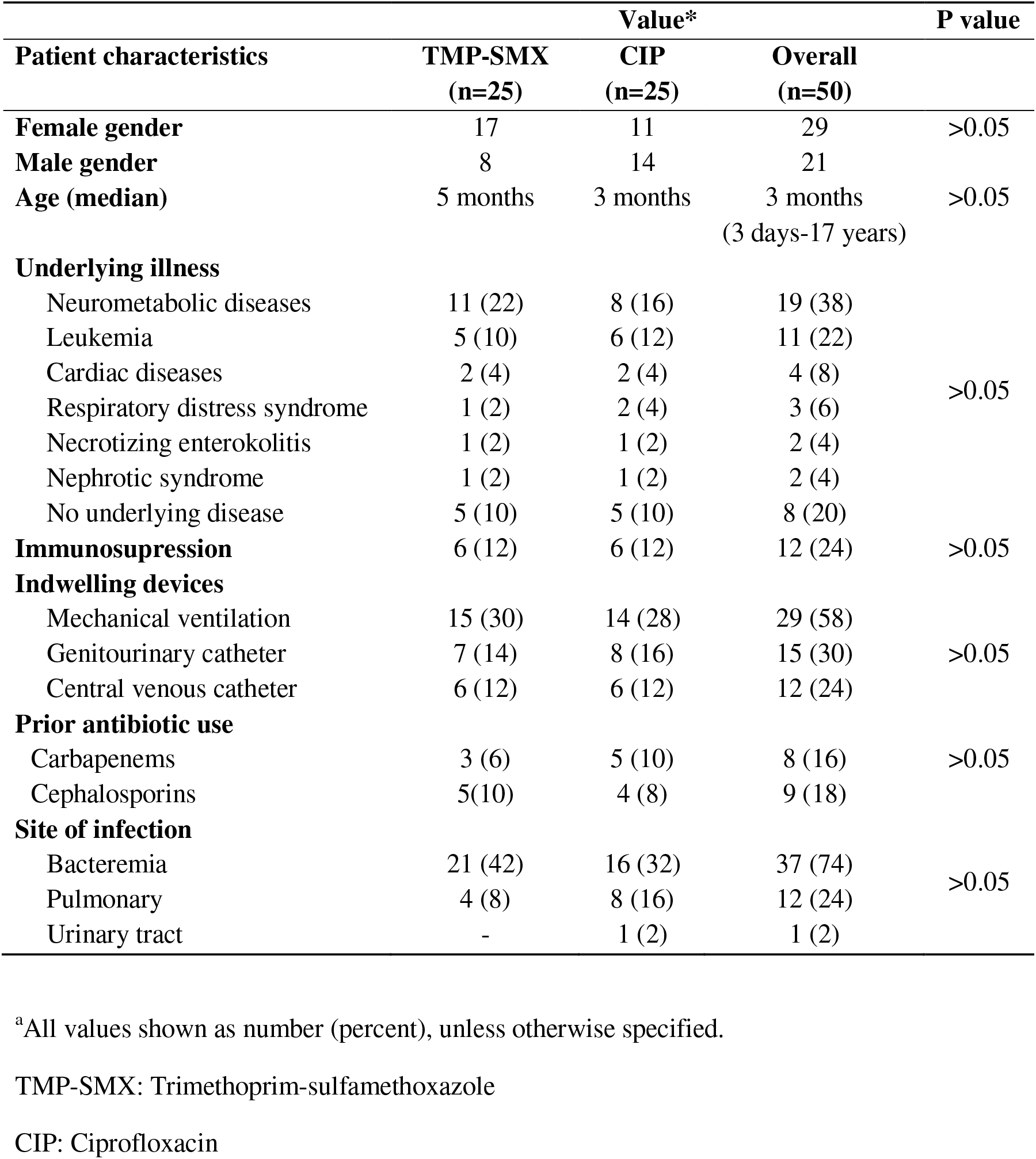
Comparison of demographic characteristics of patients with *S. maltophilia* infection who received monotherapy with trimethoprim-sulfamethoxazole or ciprofloxacin

The median duration of therapy was 11 days (IQR, 8 to 21 days) for patients who received CIP and 12 days (IQR, 3 to 21 days) for patients who received TMP-SMX *(p* =0.784) (Table 1). Eleven patients (22%) had abnormalities in laboratory tests during treatment of CIP or TMP-SMX. Four patients had side-effects in CIP group (16%) while 7 patients (28%) had side-effects in TMP-SMX group, however no significant difference was present between these groups (p>0.05). Thrombocytopenia was observed at 2 patients (4%) under CIP treatment and 3 patients (6%) under TMP-SMX. Hyponatremia developed in 1 patient (2%) at CIP group and 1 patient (2%) at TMP-SMX group. One patient at the CIP group had hypokalemia who required treatment modification. Three patients (6%) under TMP-SMX treatment had elevated alanine aminotransferase (ALT) and aspartate aminotransferase (AST) levels (Table 2).

**Table-2:** Comparison of side-effetcs of trimethoprim-sulfamethoxazole or ciprofloxacin monotherapy in children

|  | **TMP-SMX\*** | **CIP\*** | **Overall\*** |
| --- | --- | --- | --- |
| **Daystotreatment (days) [median (IQR)]** | 12 | 11 | 11.5 |
| **Abnormalities in laboratory tests** |  |  |  |
| Thrombocytopenia | 3 (6) | 2 (4) | 5 (10) |
| Hyponatremia | 1 (2) | 1 (2) | 2 (4) |
| Hypokalemia | - | 1 (2) | 1 (2) |
| Elevated liver enzymes | 3 (6) | - | 3 (6) |
| **30-days case fatality rate** | 1 (2) | 1 (2) | 2 (4) |
| **In-hospital case fatality rate** | 3 (6) | 1 (2) | 4 (8) |

\*All values shown as number (percent), unless otherwise specified.
TMP-SMX: Trimethoprim-sulfamethoxazole
CIP: Ciprofloxacin

None of the patients had died under CIP or TMP-SMX treatment following 7 days after treatment. There was no statistically significant difference between CIP and TMP-SMX groups by means of total number of side-effects (p>0.05). Thirty days and in-hospital mortality between CIP and TMP-SMX groups were not significantly different (p>0.05) (Table-2). All patients except one who had side-effects due to the both of the drugs were hospitalized at the intensive care units including 5 premature patients in NICU, 3 patients in PICU and 2 patients in cardio-vascular ICU. Among the neonates treated with TMP-SMX; the ratio of side-effects was 30% (2 of 10 patients) and was 16.7% (2 of 12 patients) with CIP, although no significant difference was present (p>0.05). The side-effect was present in 1 of 4 patients in CIP group (25%) and 2 of 5 patients (40%) in TMP-SMX group in PICU. The rate of side-effects in patients hospitalized in other wards than ICU was very low and only one patient (1 of 8 patients treated with TMP-SMX) had side-effect with the treatment.

## DISCUSSION

*S. maltophilia* is an opportunistic pathogen colonizing patients in intensive care settings, especially those with underlying debilitating conditions such as immunosuppression, malignancies, and implantation of foreign devices (catheters, respiratory therapy equipment, etc.) (3). In this study, 72% of patients were treated in the ICU. All of the patients had at least one indwelling devices and approximately 25% patients had immunosuppression.

The treatment of *S. maltophilia* is problematic because of the common multidrug resistance (MDR) of the strains. The few available antibiotics that are naturally active against *S. maltophilia* include cotrimoxazole, some β-lactams (principally the combination ticarcillin-clavulanic acid), and fluoroquinolones (essentially ciprofloxacin) (3). However the studies in children were limited.

Several studies have recommended the consideration of the use of TMP-SMX as the initial choice of the treatment of serious *S. maltophilia* infections (20,21). Fluoroquinolones, rapidly gaining prominence in treatment of *S. maltophilia*, are noted for their potency and tolerability. The previous studies showed that CIP could use alone or combination with TMP-SMX for treatment of *S. maltophilia* infections. Rojas et al. reported a case of *S. maltophilia* meningitis in a baby boy after a neurosurgical procedure, successfully treated with the combination of trimethoprim-sulfamethoxazole and ciprofloxacin (22). Lo et al. reported a preterm infant with multi-resistant *S. maltophilia* (including resistance to TMP-SMX) meningitis successfully treated with ciprofloxacin (23). Wang et al. had evaluated 98 adult patients with *S. maltophilia* infections and assessed the effectiveness of treatment with FQ monotherapy compared to TMP-SMX monotherapy (24). The outcome that patients receiving CIP showed clinical outcomes similar to those receiving TMP-SMX for the treatment of *S. maltophilia* bacteremia.

*Stenotrophomonas maltophilia* infections are associated with high morbidity and mortality, with estimated crude mortality rates ranging from 20 to 70% and with the risk of mortality highest among patients receiving inappropriate initial antimicrobial therapy (5,9,13,15,25). In our study, the 30-days mortality rate and in-hospital case fatality rate were 4% and 8% respectively. The mortality associated with *S. maltophilia* infections had been reported with a wide range between 14 to 62% (6–13,17). The variability of the mortality rates could be due to the variability of the patients and type of infections (24). Wang et al. had identified in-hospital mortality rate of 24% for all patients with *S. maltophilia* infections treated with TMP-SMX or FQ monotherapy (24). However the study group included geriatric patients with a mean age of 73 years. One report had reported all-cause mortality as low as 14 % and infection-related mortality rate of only 4% (15). The authors in the latter article suggested that high rates of patient ratio in other wards instead of ICU patients might be a factor for relatively low mortality in that study (15). However all these studies were adult studies and limited information for mortality rates in children following *S. maltophilia* infections was present with different rates. While an one study from Turkey, mortality was reported 2 of 33 (6%) in children with *S. maltophilia* infections (26); another study from Taiwan had reported the crude mortality as high as 40.6% (27). Our study had low mortality rated as 4% despite 72% of the patients were ICU patients showing the variability of mortality rates. The rate of mortality was not different between CIP and TMP-SMX group, supporting the previous studies in adult patients.

In our study; side-effects ratio were not statistically different between CIP and TMP-SMX group, however elevated liver enzymes were more prominent in the TMP-SMX group. TMP-SMX is worth mentioning especially in neonates, since previous clinical studies had demonstrated occurence of kernicterus with a sulfonamide “sulfisoxazole”, however a review of the literature showed no incidence of kernicterus in neonates (28) as in our study. However other side-effects were prominent in especially premature infants although no significant difference was present compared to CIP. Beyond the neonatal group, both drugs appeared to be good choice for *S. maltophilia* infections.

The general use of FQs in pediatric patients are restricted depending on the preclinical studies of quinolones in juvenile beagle dogs in which articular cartilage damage in weight-bearing joints(29–31) were observed and emerging concern for the occurrence of similar adverse effects in growing children. In a prospective multicenter, observational, cohort study comparing adverse events in 276 pediatric patients who had treated with FQ, adverse musculoskeletal events were reported to be more frequent comparing to the control group however no severe or persistent musculoskeletal injuries were observed at follow-up in this study (32). In a systematic review of ciprofloxacin safety in 16,184 children from 105 studies, 258 musculoskeletal adverse events was observed in 232 pediatric patients (estimated risk, 1.6; 95% CI, 0.9 to 2.6), approximately 1 musculoskeletal adverse event for every 62.5 patients. Arthralgia was the most common reported adverse musculoskeletal event (50%), affecting mostly the knee joint (33). Furthermore data from pooled safety studies; revealed was a 57% increased risk of arthropathy in the patients receiving ciprofloxacin compared to that in patients in the control arm (25). However, in another review of the literature analysis of the short-term and long-term effects of ciprofloxacin on cartilage and growth indicated no significant differences between ciprofloxacin and control groups (34). Despite many previous studies showing no significant increase in musculoskeletal complications in these children, there are still concerns about potential musculoskeletal adverse effects in young children treated with FQs.

The present study has several limitations. Firstly, treatment groups were assigned retrospectively and were not randomized. Secondly, in our study *S. maltophilia* strains were susceptible to only TMP-SMX and CIP; so we found the mortality rate lower than other studies. Therefore, our results could not be generalized for multidrug-resistant *S. maltophilia* strains. In the previous studies, major limitation reported was failure to to distinguish between colonization and infection with *S. maltophilia* may also influence clinical success (15). However we could overcome this limitation by including only patients with documented bacteremia with *S. maltophilia.*

## CONCLUSION

CIP had the same clinical outcome and mortality as TMP-SMX for the treatment of *S. maltophilia* bacteremia. Our data suggest that either CIP or TMP-SMX would be appropriate choice in for isolates known to be susceptible. Due to the feared arthrotoxicity associated with FQ, CIP should be considered in cases in which TMP-SMX could not be used or stopped due to the side-effects. Further prospective randomized studies are required for comparing CIP and TMP-SMX for the treatment of *S. maltophilia* infections in pediatric patients including neonates.

## Acknowledgement

The Authors declare that there is no conflict of interests.

## Author Contributions

**Ahu Aksay**: Concept, Design, Data Collection and/or Processing, Literature review, Writing.

**İlker Devrim**: Concept, Design, Analysis and/or Interpretation, Supervision, Literature review, Writing, Critical Review.

**Nuri Bayram**: Analysis and/or Interpretation, Supervision, Critical Review, Literature review.

**Nagehan Katipoglu**: Literature review.

**Gökhan Ceylan**: Literature review.

**Şebnem Çalkavur**: Literature review, Supervision.

**Sertaç Arslanoglu**: Literature review, Supervision.

**Hasan Agın**: Analysis and/or Interpretation, Supervision, Critical Review.

**Gamze Gulfidan**: Data Collection and/or Processing, Literature review, Writing.

